# Haplotype-phased synthetic long reads from short-read sequencing

**DOI:** 10.1101/022897

**Authors:** James A. Stapleton, Jeongwoon Kim, John P. Hamilton, Ming Wu, Luiz C. Irber, Rohan Maddamsetti, Bryan Briney, Linsey Newton, Dennis R. Burton, C. Titus Brown, Christina Chan, C. Robin Buell, Timothy A. Whitehead

**Affiliations:** Department of Chemical Engineering and Materials Science, Michigan State University, East Lansing, MI; Department of Plant Biology, Michigan State University, East Lansing, MI; Michigan Center for Translational Pathology, University of Michigan, Ann Arbor, 48109, USA; Department of Microbiology and Molecular Genetics and Department of Computer Science and Engineering, Michigan State University, East Lansing, MI; BEACON Center for the Study of Evolution in Action, Michigan State University, East Lansing, MI; Department of Integrative Biology, Michigan State University, East Lansing, MI; Department of Immunology and Microbial Science, The Scripps Research Institute, La Jolla, CA; Center for HIV/AIDS Vaccine Immunology and Immunogen Discovery, The Scripps Research Institute, La Jolla, CA; International AIDS Vaccine Initiative Neutralizing Antibody Center, The Scripps Research Institute, La Jolla, CA; Population Health and Reproduction, University of California, Davis, CA; Department of Biosystems and Agricultural Engineering, Michigan State University, East Lansing, MI

**Keywords:** synthetic long reads, DNA sequencing

## Abstract

Next-generation DNA sequencing has revolutionized the study of biology. However, the short read lengths of the dominant instruments complicate assembly of complex genomes and haplotype phasing of mixtures of similar sequences. Here we demonstrate a method to reconstruct the sequences of individual nucleic acid molecules up to 11.6 kilobases in length from short (150-bp) reads. We show that our method can construct 99.97%-accurate synthetic reads from bacterial, plant, and animal genomic samples, full-length mRNA sequences from human cancer cell lines, and individual HIV *env* gene variants from a mixture. The preparation of multiple samples can be multiplexed into a single tube, further reducing effort and cost relative to competing approaches. Our approach generates sequencing libraries in three days from less than one microgram of DNA in a single-tube format without custom equipment or specialized expertise.

## Introduction

The short-read assembly paradigm currently dominates genomics (Margulies et al. 2005; Bentley et al. 2008). However, the loss of linkage information during the generation of short reads limits their utility. In particular, short reads are insufficient to phase the haplotypes of individuals within mixtures of similar sequences, including polyploid chromosomes (Consortium et al. 2011; Jia et al. 2013), viral quasispecies (Acevedo et al. 2014), multiply or alternatively spliced mRNA (Menon et al. 2014), genes from metagenomic samples containing related organisms (Hess et al. 2011; Sharon et al. 2015), and immune antibody gene repertoires (Georgiou et al. 2014). In these cases, additional information is required to determine whether mutations separated by distances longer than the read length are present in the same individual.

This limitation can be overcome by technologies that provide long continuous sequence reads. “True” long read technologies (Branton et al. 2008; Metzker 2010) hold great promise, but currently suffer from high error rates. As an alternative, sample preparation protocols that enable long “synthetic” reads to be constructed from conventional short reads (Miller et al. 2007; Hiatt et al. 2010; Lundin et al. 2013; Voskoboynik et al. 2013; Hong et al. 2014; Kuleshov et al. 2014; McCoy et al. 2014; Wu et al. 2014). However, these approaches have been limited by generalizability, cost, throughput, or synthetic read length.

Here, we present a library preparation method that overcomes these limitations, providing a general platform for affordable and reliable synthetic read generation from a wide range of input nucleic acid types. In contrast to competing approaches, our method requires no specialized equipment or proprietary software and is carried out in a single tube. We demonstrate generalizability by assembling synthetic reads from a range of samples including genomic DNA from bacteria, plants, and an animal, mRNA isolated from human cancer lines, and mock viral patient samples. We validate 99.97%-accurate synthetic reads up to 11.6 kb in length, and show their utility for improving a plant draft genome assembly. We show that the preparation of multiple samples can be multiplexed into a single tube, further reducing effort and cost relative to competing approaches. Finally, using synthetic reads, we directly observe up to thirty-five splice junctions in an individual mRNA and individual haplotypes from mixtures of highly similar molecules with a low rate of chimera formation.

## Results

The approach is illustrated schematically in Figure 1A. DNA fragments of lengths up to 20 kb are appended at each end with adapters containing a degenerate barcode region flanked by defined sequences, such that every target fragment becomes associated with two unique barcodes. PCR with a single primer produces many copies of each target molecule along with its two associated barcodes. The priming sites are removed and a single break (on average) is enzymatically induced in each copy to expose regions of unknown sequence at the newly created ends. Those regions are brought into proximity with the barcode at the opposite end of each fragment by intramolecular circularization (Fig. S1). Next, the molecules are linearized and a sequencing-ready library compatible with the Illumina platform is prepared (Supplementary Fig. 2). The resulting short reads begin with the barcode sequence and continue into the unknown region. Following sequencing, reads are grouped by common barcodes. The two distinct barcodes appended to each target molecule can be identified and paired (Fig. 1A, Supplementary Fig. 3), allowing the two barcode-defined groups derived from the two ends of the original fragment to be combined. The read groups are assembled independently and in parallel to reconstruct the full sequences of the original DNA molecules.

**Figure 1.**
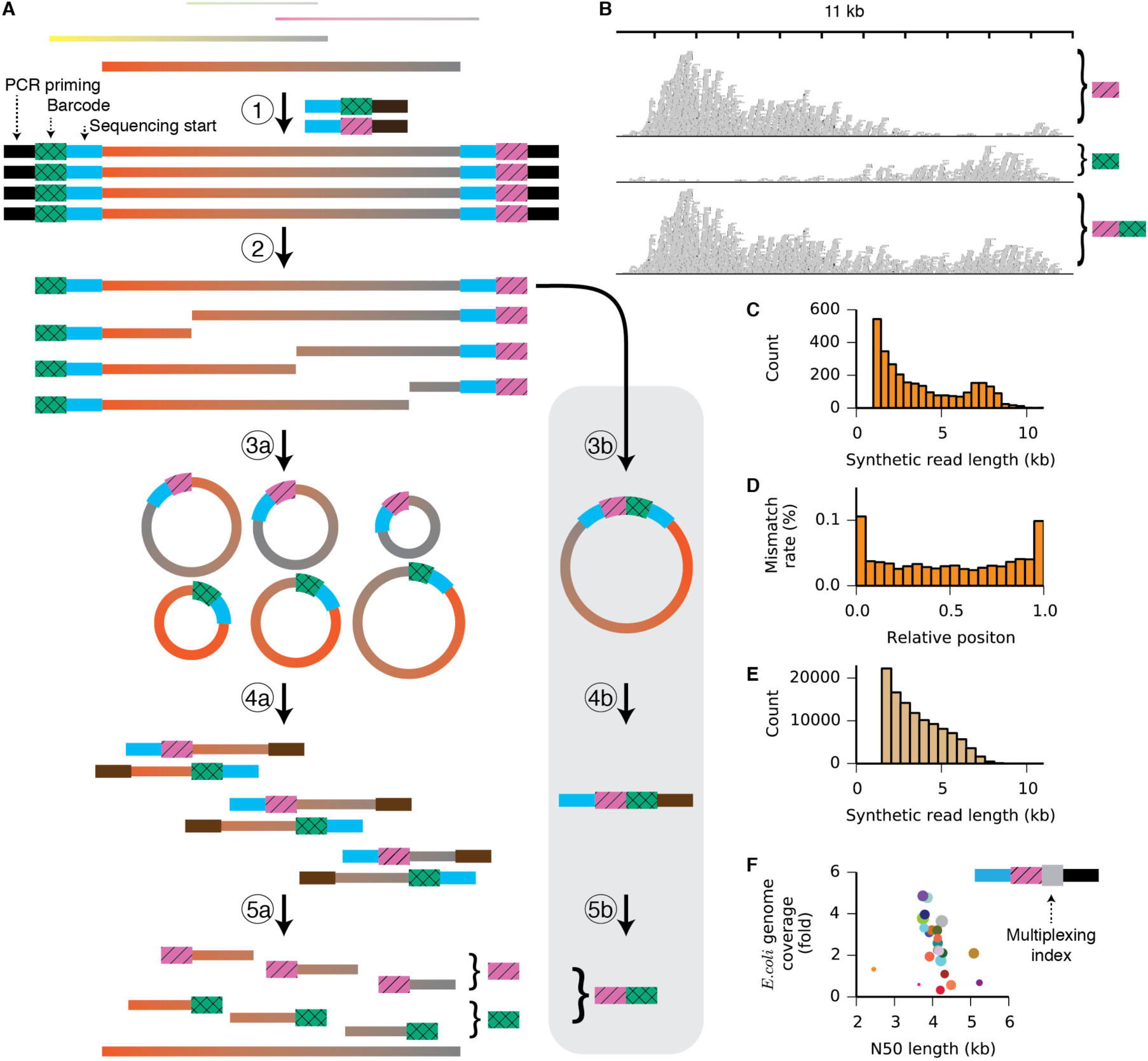
A method for assembling synthetic long reads. *(A)* Schematic of the approach. A supplemental barcode-pairing protocol (grey box) resolves the two distinct barcodes affixed to each original target molecule. *(B)* Reads associated with two distinct barcodes are shown aligned to the *E. coli* MG1655 reference genome. Barcode pairing merges the groups (bottom), increasing and evening the coverage and allowing assembly of the full 10-kb target sequence. *(C)* Length histogram of synthetic reads assembled from *E. coli* MG1655 genomic reads (minimum length 1 kb). The N50 length of the synthetic reads is 6.0 kb, and the longest synthetic read is 11.6 kb. *(D)* Mismatch rates of synthetic reads from the *E. coli* MG1655 dataset as a function of relative position along the synthetic read. *(E)* Length histogram of synthetic long reads assembled from *Gelsemium sempervirens* genomic reads (minimum length 1.5 kb). The N50 length of the synthetic reads is 4.3 kb. *(F)* An additional multiplexing index region (grey square) allows adapter-ligated samples to be mixed and processed in a single tube. Genomic DNA from twenty-four experimentally evolved strains of *E. coli* was separately ligated to adapters and amplified, then mixed into a single tube for the remaining steps of the protocol. *E. coli* genome coverage and N50 length are plotted for synthetic reads from each strain. Circle size indicates the number of short reads demultiplexed to a given strain.

To validate the accuracy of the synthetic long reads, we prepared sequencing libraries from genomic DNA isolated from *E. coli* MG1655. Illumina 150-bp paired-end reads were trimmed to remove barcodes, adapter sequences, and regions of low quality, and sorted into barcode-delineated groups. When aligned to the MG1655 reference genome, more than 80% of the reads in each barcode group aligned to the same 10-kb region. The coverage distribution across the region was non-uniform, dropping off with distance from the barcoded end (Fig. 1B). A total of 1,215 barcode pairs were identified and their read groups merged. Because coverage from one barcode is high in the region of the target molecule where coverage from its partner is low, merging groups not only increases but also evens the coverage across the target. After barcode pairing, 2,792 read groups contained at least fifty read pairs. Independent *de novo* assembly of each group yielded 2,878 synthetic reads of length greater than 1 kb (Fig. 1C, Supplementary Table 1), with the longest reaching 11.6 kb. Barcode pairing improved the N50 assembly length and reduced the number of redundant synthetic reads (Supplementary Fig. 4 and 5). To determine the fidelity of assembly, synthetic reads longer than 1.5 kb were aligned to the MG1655 reference genome (Hayashi et al. 2006). The mismatch rate within the aligned regions of the synthetic reads was 0.04% (Supplementary Table 2). Errors were more common at the ends of the synthetic reads, where short-read coverage was low (Fig. 1D and Supplementary Fig. 6). When 100 nucleotides were trimmed from each end of the synthetic reads, the mismatch rate dropped to 0.03%. Transition mutations (C/T and A/G) made up 75.6% of all mismatches, suggesting that the PCR amplification step is the dominant source of error (Dunning et al. 1988).

We further evaluated our method with genomic DNA isolated from higher organisms with well-developed draft genome assemblies. From *Gallus gallus* (chicken) genomic DNA (Rubin et al. 2010), we assembled 125,203 synthetic reads longer than 1 kb, with an N50 length of 2.0 kb. The length distribution (Supplementary Fig. 7) and low N50 length relative to the shearing length indicate that this library was under-sequenced, and additional sequencing would yield longer synthetic reads. Nonetheless, all but 661 (0.13%) of the 510,070 synthetic reads of all lengths aligned to the *G. gallus* reference genome. Additionally, we generated 1,411 synthetic reads longer than 1 kb (N50 of 3.1 kb) from a doubled-monoploid potato (*Solanum tuberosum* Group Phureja) (Consortium et al. 2011; Sharma et al. 2013) (Supplementary Fig. 7). 97.3% of the synthetic reads aligned to the draft reference genome (Supplementary Table 3).

Having validated the method, we used synthetic long reads to improve a shotgun assembly of the genome of *Gelsemium sempervirens,* an ornamental and medicinal plant species with an estimated genome size of 312 Mb. A draft assembly was first created from 161.4 million Illumina whole-genome shotgun reads. We then prepared a barcoded library and assembled 111,054 synthetic reads longer than 1.5 kb (Fig. 1E and Supplementary Fig. 8), with an assembly N50 length of 4.3 kb (Supplementary Table 1). A total of 397.8 Mb of synthetic reads (Supplementary Table 4) were used to scaffold the assembly. Incorporation of synthetic reads improved the assembly from 25,276 contigs with an N50 contig length of 19,656 bp to 18,106 scaffolds with an N50 scaffold length of 29,078 bp (Table 1). Maximum contig length also increased, from 198 kb in the shotgun assembly to 366 kb in the synthetic read assembly. Assembly quality metrics generated with the CEGMA pipeline (Supplementary Table 5) and by alignment of cleaned RNA-seq reads (Supplementary Table 6) were consistent with an increased representation of the genome after incorporation of the synthetic long reads.

**Table 1.** Genome assembly statistics for *G. sempervirens*.

|  | Shotgun contigs | Synthetic read scaffolds |
| --- | --- | --- |
| Contig/scaffold N50 size (bp) | 19,656 | 29,078 |
| Total assembly size (bp) | 215,038,998 | 218,719,799 |
| No. of contigs/scaffolds | 25,276 | 18,106 |
| Maximum length (bp) | 197,779 | 365,589 |
| Small contigs (< 1 kb) were filtered out. |  |  |

In contrast to dilution-based synthetic read approaches (Voskoboynik et al. 2013; Kuleshov et al. 2014; McCoy et al. 2014), the intramolecular circularization step in our protocol makes it possible to prepare a library in a single tube. We further exploited this property to allow multiple samples to be combined and prepared in the same mixture, yielding considerable savings in cost and time. To accomplish this, we modified the adapters to include multiplexing index sequences between the PCR priming region and the molecule-specific barcode (Fig. 1f, Supplementary Fig. 9). Genomic DNA from each of twenty-four laboratory-evolved *E. coli* strains (Souza V et al. 1997) was isolated, sheared to 6-10 kb, ligated to adapters containing both a multiplexing index unique to each strain and a molecule-specific barcode, and amplified by PCR. Purified PCR products were then mixed and the remainder of the library preparation protocol was performed on the single mixed sample. Sequencing reads were demultiplexed by project according to standard 6-bp index read, then further demultiplexed by strain according to the barcode-adjacent multiplexing index identified in the forward read, sorted by barcode, and assembled in parallel (Supplementary Fig. 10, Supplementary Table 7). The summed lengths of the synthetic reads longer than 1 kb exceeded twofold genome coverage for sixteen out of the twenty-four strains, with a median genome coverage of 2.3 fold and median N50 of 4.1 kb. Cases of low coverage resulted from either adding insufficient amplified DNA into the multiplexed mixture (small circles, Fig. 1F) or carrying too few doubly barcoded molecules into the amplification step (large circles with high N50 but low coverage, Fig. 1F). Along with the nanogram-level input requirements, single-tube format, and competitive assembly length metrics demonstrated earlier, single-tube multiplexing provides a substantial advantage in throughput, convenience, and cost over competing synthetic read techniques for genome assembly and phasing (Supplementary Table 8).

We next asked whether our method could be extended to applications beyond genome assembly and phasing. In RNA-seq experiments, the presence of transcripts resulting from multiple splicing events must be inferred statistically because individual reads are too short to regularly span multiple splice junctions. We used synthetic long reads to directly observe multiply spliced messenger RNA isolated from human cancer cells. We modified the Smart-seq2 method (Picelli et al. 2013) to incorporate adapters with molecule-specific barcodes during the reverse transcription and second-strand synthesis steps (Supplementary Fig. 11). The barcoded cDNA product was amplified, broken, circularized, and prepared for sequencing. From mRNA isolated from HCT116 and HepG2 cells, we assembled 28,689 and 16,929 synthetic reads, respectively, of lengths between 0.5 and 4.6 kb (Fig. 2). Approximately 97% of the splice junctions captured by the synthetic long reads have been observed previously (Fig. 2C, Supplementary Tables 9-11). In contrast to conventional RNA-seq reads, synthetic reads spanned multiple splice junctions (Fig. 2C, Supplementary Fig. 12), with a median of 2.0 spanned junctions per synthetic read for both samples and a maximum of 35 spanned junctions. Examination of the synthetic reads revealed examples of differential splicing between the HCT116 and HepG2 cell lines, as well as a novel transcript in the HCT116 cell line (Supplementary Fig. 13). Notably, our protocol can be adapted for single-cell RNA-seq methods, potentially allowing discrimination of multiple splice variants for individual cells.

**Figure 2.**
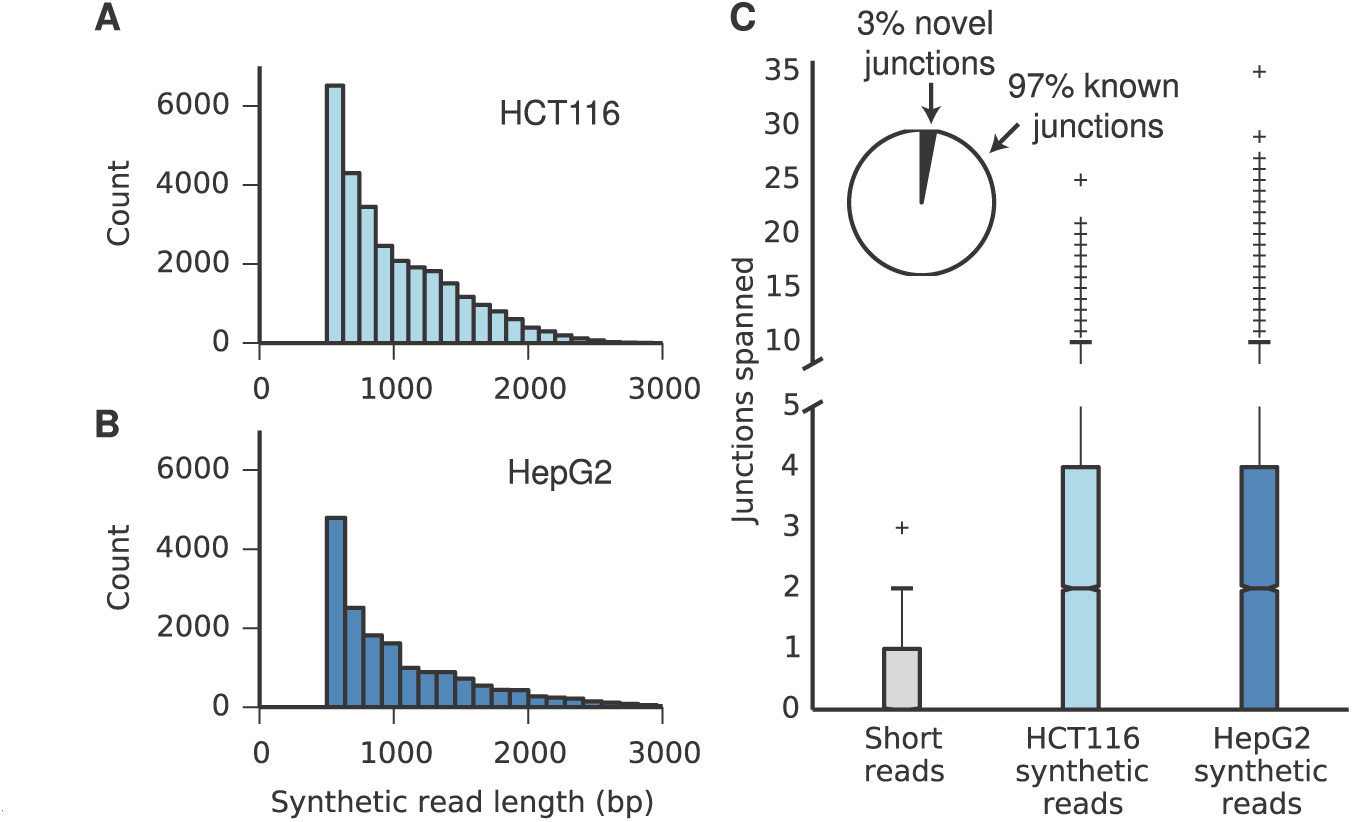
Individual assembly of full-length mRNA sequences. *(A)* Length distribution of synthetic long reads (minimum length 500 bp) from HCT116 mRNA. *(B)* Length distribution of synthetic long reads (minimum length 500 bp) from HepG2 mRNA. *(C)* Box plots showing the number of splice junctions spanned by short reads and synthetic long reads. The axis is broken between 5-10 junctions spanned and the scale changed; a version with a standard axis is presented as Supplementary Figure 12. Inset: 97% of the junctions identified in the synthetic reads are known, providing validation for the method.

Highly accurate haplotype-phased sequences open new routes to studying mixtures of similar yet distinct individuals. These include viral quasispecies, which consist of near-identical genomes harboring key mutations, isolated B-cells, whose genomes encode therapeutic antibodies differing mainly in a few hypervariable regions, and environmental samples, which contain closely related strains with homologous genes and operons. In particular, the ability to generate thousands of complete, individual, haplotype-phased viral genome sequences would provide an unprecedented view of the ways viruses adapt to immune responses and environmental changes such as the introduction of antiretroviral therapies. The TruSeq Synthetic Long Reads approach is incompatible with mixtures of similar molecules because it assigns a barcode to a diluted pool of a few hundred target molecules. When drawn from a large genome these molecules are unlikely to overlap, but when all molecules are similar each must be diluted into its own well, limiting throughput.

First, to demonstrate the ability to resolve and haplotype individual genes, we generated barcoded reads from a mixture of plasmids. As expected, each barcode-defined group of reads was dominated by sequences unique to a single parent (Supplementary Fig. 14). Next, we demonstrated viral haplotyping by sequencing a mixture of two *env* gene variants. *Env* is an HIV gene that encodes the envelope glycoprotein, which triggers infection by binding CD4 and a co-receptor (either CXCR4 or CCR5) and is the primary target for HIV vaccine development (Burton et al. 2004). A large fraction of the *de novo*-assembled synthetic long reads approached the expected full length of 3 kb (Fig. 3A). We were able to unambiguously identify 1,157 out of 1,173 (98.6%) assembled and cleaned sequences as one or the other of the two variants. The sixteen outlier reads that do not map to one of the two clusters (Fig. 3C) have high mismatch rates against both parent sequences, rather than intermediate mismatch with both as would be expected of chimeric sequences. These results indicate accurate assembly and minimal chimera formation, and we conclude that the method is suitable to detect rare individual viral genomes harboring distant interacting loci.

**Figure 3.**
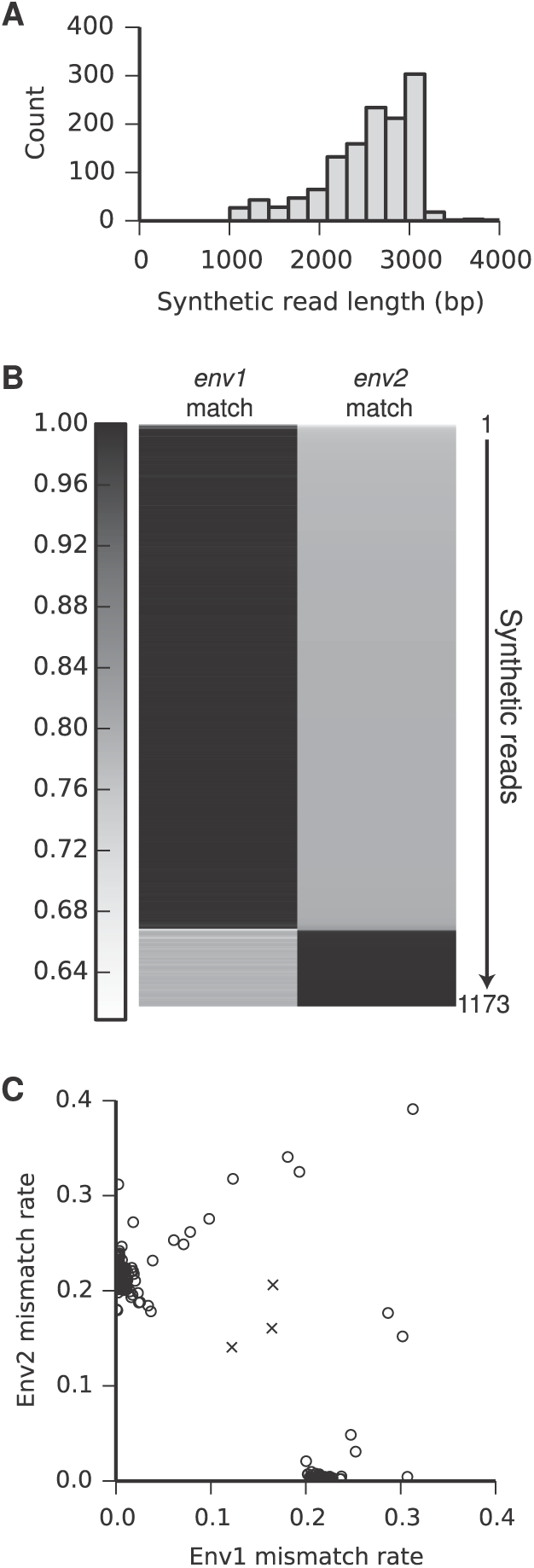
Individual assembly of full-length *env* genes from a mixture of two variants. *(A)* The length distribution of the synthetic long reads (minimum length 1 kb) shows assembly of full-length 3-kb *env* gene sequences. *(B)* 1,173 synthetic reads between 1.5 and 3.2 kb in length were aligned to each of the two original *env* sequences (*env1* and *env2*). The alignment match rates are shown as a heatmap, with each synthetic read represented by a thin horizontal line. The majority of the synthetic reads align with low error to exactly one of the two original sequences, indicating high accuracy and a low rate of chimera formation. Chimeric reads would be expected to match both original sequences at intermediate accuracies. *(C)* Scatter plot showing the mismatch rates of each synthetic read against the two known *env* sequences. Synthetic reads (open circles to emphasize extensive overlap) cluster into two distinct groups along the axes (near-zero mismatch rate). Even the sixteen reads that do not fall on the clusters are distant from three manually created mock chimeras (crosses), indicating a low frequency of chimera formation.

## Discussion

We have demonstrated a versatile approach to assemble individual DNA sequences from mixed-population short-read data from a variety of DNA and RNA samples. Our approach offers important advantages over competing synthetic read methods (Supplementary Tables 8 and 12). Many samples can be multiplexed and prepared for sequencing in three days in a single tube with no custom equipment or specialized expertise, providing significant benefits in cost and throughput. Our method can be applied to a wider range of sample types than competing technologies, from genomic fragments to uniform mixed populations, and its novel barcode pairing protocol yields longer synthetic reads than previous circularization-based approaches.

Because synthetic reads are built from overlapping short reads, improved read length and accuracy come at a cost of sequencing depth. In the examples presented here, between 43 and 225 nucleotides of short-read sequencing were used to generate each nucleotide of synthetic read (Supplementary Table 1). The cost of synthetic long reads is justified by additional linkage information, improved accuracy, and lowered resource demand through parallelization of the computational pipeline. Synthetic reads will become increasingly attractive as sequencing throughput increases and costs drop, further justifying the exchange of quantity for quality.

## Materials and methods

### Library preparation for synthetic long read assembly

A 100 μL solution of two oligonucleotides (oligos 1 and 2, Supplementary Table 13, at 2 uM and 5 uM, respectively) in NEB buffer 2 (New England Biolabs, Ipswich, MA) was heated to 95°C for 10 minutes and allowed to cool slowly to 37°C. 5 units of Klenow exo- (NEB) and 0.3 mM each dNTP (NEB) were added and the mixture was incubated at 37°C for 60 minutes. The DNA to be sequenced was typically diluted to 50 μL at 10 ng/μL and fragmented into ∼10 kb pieces with a g-TUBE (Covaris, Woburn, MA) by centrifugation at 4200 g according to the manufacturer’s protocol. The DNA was end-repaired with the NEBNext End Repair Module (NEB) according to the manufacturer’s protocol, purified with a Zymo column (Zymo Research, Irvine, CA) and eluted in 8 μL buffer EB (Qiagen, Hilden, Germany). The DNA was then dT-tailed by incubation in 1X NEB buffer 2 with 1 mM dTTP (Life Technologies, Carlsbad, CA), 5 units Klenow exo-, and 10 units polynucleotide kinase (NEB) at 37°C for 1 hour. 250 fmol of library DNA (4 μL) and 5 pmol of barcode adapters (1 μL) were ligated with 5 μL TA/Blunt MasterMix (NEB) according to the manufacturer’s protocol, gel purified with the Qiagen Gel Extraction kit if necessary, and eluted in 50 μL buffer EB. The concentration of doubly ligated product was determined by qPCR and/or dilution-series PCR. The library was diluted to about 100,000 doubly ligated molecules per μL. Approximately 100,000 molecules of adapter-ligated DNA were amplified by PCR in 150 μL reactions split across eight PCR tubes with LongAmp Taq DNA polymerase (NEB) using a single primer (oligo 2, Supplementary Table 13) (Rungpragayphan et al. 2002; Stapleton and Swartz 2010) at 0.5 mM and the following thermocycling conditions: 92°C for 2 minutes, followed by 40 cycles of 92°C for 10 seconds, 55°C for 30 seconds, and 65°C for 3 minutes/kb, followed by a final hold at 65°C for 10 minutes.

The PCR product was gel purified with the Qiagen Gel Extraction kit, eluted in 50 μL of buffer EB, and quantified by absorbance at 260 nm. 200 ng to 1 μg of DNA were mixed with 1 unit of USER enzyme (NEB) in a 45 μL reaction volume and incubated for 30 minutes at 37°C. 100 ug/mL bovine serum albumin and 5 μL of dsDNA fragmentase buffer were added and the mixture incubated on ice for 5 minutes. 0.5-2 μL of dsDNA fragmentase (NEB) (volume adjusted based on amount and length of DNA to be fragmented) were added and the mixture was incubated at 37°C for 15 minutes. The reaction was stopped by addition of 5 μL of 0.5 M EDTA and fragmentation was confirmed by the presence of a smear on an agarose gel. The DNA was purified with 0.8 volumes of Ampure XP beads (Beckman Coulter, Brea, CA), eluted in 20 μL buffer EB, and quantified by absorbance at 260 nm. 2 μL of 10X NEB Buffer 2 were added and fragmented DNA was incubated with 0.5 μL of *E. coli* DNA ligase (NEB) for 20 minutes at 20°C. 3 units T4 DNA polymerase, 5 units Klenow fragment (both from NEB), and 50 μM biotin-dCTP (Life Technologies) were added and the reaction was incubated for 10 minutes at 20°C. 50 μM dGTP, dTTP, and dATP were added and the mixture was incubated for an additional 15 minutes, purified with 1 volume of Ampure XP beads, eluted in 20 μL buffer EB, and quantified by absorbance at 260 nm. 200-1000 ng of DNA at a final concentration of 1 ng/μL were mixed with 3000 units of T4 DNA ligase and T4 DNA ligase buffer to 1X, and incubated at room temperature for 16-48 hours.

Linear DNA was digested by addition of 10 units of T5 exonuclease and incubation at 37°C for 60 minutes. Circularized DNA was purified with a Zymo column and eluted in 130 μL buffer EB. The DNA was fragmented with the Covaris S2 disruptor to lengths ∼500 bp. 20 μL of Dynabeads M-280 Streptavidin Magnetic Beads (Life Technologies) were washed twice with 200 μL of 2X B&W buffer (1X B&W buffer: 5 mM Tris-HCl (pH 7.5), 0.5 mM EDTA, 1 M NaCl) and resuspended in 100 μL of 2X B&W buffer. The DNA solution was mixed with this bead solution and incubated for 15 minutes at 20°C. The beads were washed twice with 200 μL of 1X B&W buffer and twice in 200 μL of Qiagen buffer EB. At this point, 15% (30 μL) of the beads were removed to a new tube for two-tube barcode pairing (see below). The remaining beads were resuspended in NEBNext End Repair Module solution (42 μL water, 5 μL End Repair Buffer, and 2.5 μL End Repair Enzyme Mix), incubated at 20°C for 30 minutes, and washed twice with 200 μL of 1X B&W buffer and twice with 200 μL of buffer EB. The beads were then resuspended in 17 μL water, 2 μL NEB buffer 2, 0.5 μL 10 mM dATP, and 5 units Klenow exo- polymerase, incubated at 37°C for 30 minutes to add dA tails to the DNA fragments, and washed twice with 200 μL of 1X B&W buffer and twice with 200 μL of buffer EB. A 15 micromolar equimolar mixture of two oligonuleotides (oligos 3 and 4, Supplementary Table 13) in 1X T4 DNA ligase buffer was incubated at 95°C for 2 minutes and allowed to cool to room temperature over 30 minutes. The beads were resuspended in a solution consisting of 5 μL of NEB Blunt/TA ligase master mix, 0.5 μL of 15 micromolar adapter oligo solution, and 4 μL of water. The mixture was incubated for 15 minutes at room temperature. The beads were washed twice with 200 μL of 1X B&W buffer and twice with 200 μL of buffer EB. For amplification by limited-cycle PCR (lcPCR), the beads were resuspended in a 50 μL PCR solution consisting of 36 μL of water, 10 μL of 5X Phusion HF DNA polymerase buffer, 1.25 μL of each of 10 micromolar solutions of Illumina Index and Universal primers (oligos 5 and 6, Supplementary Table 13), and 0.02 units/μL Phusion DNA polymerase (Thermo Fisher Scientific, Waltham, MA). The following thermocycling program was used: 98 degrees for 30 seconds, followed by 18 cycles of 98°C for 10 seconds, 60°C for 30 seconds, and 72°C for 30 seconds, and a final hold at 72 degrees for 5 minutes. The supernatant was retained and the beads discarded. The PCR product was purified with 0.7 volumes of Ampure XP beads and eluted in 10 μL buffer EB, or 500-900 bp fragments were size-selected on an agarose gel, gel-purified with the Qiagen MinElute Gel Extraction kit, and eluted in 15 μL of buffer EB. The size distribution of the DNA was measured with an Agilent bioanalyzer and cluster-forming DNA was quantified by qPCR. The DNA fragments were sequenced on an Illumina MiSeq, NextSeq, or HiSeq with standard Illumina primer mixtures.

### Two-tube barcode pairing

See Supplementary Note and Supplementary Figure 3. Bead-bound DNA was digested with 10 units of SapI in 1X CutSmart buffer in a 20 μL total volume for 1h at 37C. The beads were washed three times with 200 μL of 1X B&W buffer and twice with 200 μL of buffer EB. A 15 μM equimolar mixture of two oligonucleotides (oligos 7 and 8, Supplementary Table 13) in 1X T4 DNA ligase buffer was incubated at 95°C for 2 minutes and allowed to cool to room temperature over 30 minutes. The beads were resuspended in a solution consisting of 5 μL of NEB Blunt/TA ligase master mix, 0.5 μL of 15 μM adapter oligo solution, and 4 μL of water. The mixture was incubated for 15 minutes at 4°C and 15 minutes at 20°C. The beads were washed twice with 200 μL of 1X B&W buffer and twice with 200 μL of buffer EB. For amplification by limited-cycle PCR, the beads were resuspended in a 50 μL PCR solution consisting of 36 μL of water, 10 μL of 5X Phusion HF DNA polymerase buffer, 1.25 μL of each of 10 μM solutions of two primers (oligos 6 and 9, Supplementary Table 13, with oligo 9 selected to have a different multiplexing index than oligo 5 used above), and 0.02 units/μL Phusion DNA polymerase (Thermo Fisher Scientific). The following thermocycling program was used: 98°C for 30 seconds, followed by 18 cycles of 98°C for 10 seconds, 60°C for 30 seconds, and 72°C for 30 seconds, and a final hold at 72°C for 5 minutes. The supernatant was retained and the beads discarded. DNA was purified with 1.8 volumes of Ampure XP beads and eluted in 10 μL buffer EB. The expected product size of ∼170 bp was confirmed by agarose gel electrophoresis and Agilent bioanalyzer. Cluster-forming DNA was quantified by qPCR. The DNA fragments were mixed with the main library so as to comprise 1-5% of the total molecules, and sequenced on an Illumina MiSeq, NextSeq, or HiSeq with standard Illumina primer mixtures.

### Single-tube barcode pairing

See Supplementary Note, Supplementary Figure 15. Two different versions of oligo 1 (Supplementary Table 13) were mixed, extended with oligo 2 (Supplementary Table 13), and ligated to dT-tailed target fragments as above. The library preparation protocol was carried out as above, except that the extra barcode-pairing steps were omitted. Limited-cycle PCR was performed with 1.25 μL of a 10 micromolar solution oligo 10 in addition to oligos 5 and 6 (Supplementary Table 13).

### Complexity determination

A key step in the protocol is the quantification of doubly barcoded fragments prior to PCR. Doubly barcoded fragment concentration in this study was estimated in three ways: quantitative PCR with a quenched fluorescent probe (probe 1, Supplementary Table 13), dilution series endpoint PCR, and quantification by next-generation sequencing. For the latter, barcoded molecules were purified and serially diluted. Four dilutions were amplified with oligo 6 and four versions of oligo 11 (Supplementary Table 13) containing different multiplexing index sequences. The resulting products were mixed and sequenced with 50-bp single-end reads on an Illumina MiSeq. Reads were demultiplexed and unique barcodes at each dilution were counted. When combined with the multiplexed library preparation strategy, which enables further demultiplexing on the basis of an index in the forward read, many samples can be quantified in a single MiSeq run.

### Library preparation for synthetic long read assembly from mRNA samples

Full-length reverse transcripts were prepared essentially as in (Picelli et al. 2014), with modified primers. The “oligo-dT primer” and “TSO primer” were replaced by oligo 12 and oligo 13 (Supplementary Table 13), respectively. Barcoded full-length reverse transcripts were then processed and sequenced as above, starting from the library quantification step.

### Mulitplexed sample preparation

Two *E. coli* strains were isolated from each of the twelve recombination treatment populations in Souza *et al*. (Souza V et al. 1997). Genomic DNA was isolated from each of the twenty-four strains, sheared, end-repaired, and dT-tailed as described above in separate tubes. Twenty-four barcode adapters (Supplementary Table 14), identical except for distinct 6-bp multiplexing index regions adjacent to the barcode sequence, were prepared and ligated to the genomic fragments as described above. Adapter-ligated DNA was PCR amplified as above. Purified PCR products were quantified and equal amounts were combined into a single mixture. This mixture was prepared for sequencing following the remaining steps of the above protocol.

### Assembly of synthetic long reads

Barcoded short reads were assembled into synthetic long reads with custom python scripts, available for download at www.github.com/jstapleton/synthetic_reads. In the first step, low-quality regions and barcode sequences are removed using Trimmomatic (v 0.30) (Bolger et al. 2014) and overlapping paired-end reads are merged with FLASh (v 1.2.10) (Magoc and Salzberg 2011). In the second step, reads are further trimmed and sorted into groups according to their barcodes. In the third step, each group is *de novo* assembled independently with the SPAdes assembler (v 3.1.1) (Bankevich et al. 2012).

### Mismatch rate calculation

Synthetic long reads 1500 bp in length or longer assembled from *E. coli* K12 MG1655 genomic sequencing reads were aligned to the reference genome^20^ using BWA-MEM (v 0.7.12)^33^. The resulting SAM file was parsed with a custom script to extract mismatch, insertion, deletion, and clipping frequencies.

### Alignment of synthetic reads to the *S. tuberosum* assembly

Potato synthetic reads were trimmed by removing 100 nt from each end and then removing reads shorter than 1 kb. The trimmed synthetic reads were then aligned to the *S. tuberosum* Group Phureja DM1-3 pseudomolecules (v 4.03) (Consortium et al. 2011; Sharma et al. 2013) with BWA-MEM (v 0.7.12) (Li 2013). A custom Perl script was used to bin the alignments based on the mapping quality score for the alignment.

### Genome assembly of *Gelsemium sempervirens*

DNA was isolated from *G. sempervirens* (“Carolina Jessamine”) using the CTAB method (Saghai-Maroof et al. 1984). DNA was sheared to 300, 500, and 700 bp using a Covaris S2 instrument, end repaired using NEBNext End Repair Module (New England Biolabs, Ipswich, MA) and dA-tailed using Klenow Fragment (New England Biolabs) prior to ligation with annealed universal adaptors. The ligated DNA fragments were size selected using the Agencourt AmPure XP beads (Beckman Coulter, Indianapolis, IN), and then amplified with indexed primers for eight cycles with HiFi HotStart DNA polymerase (KAPA Biosystems, Wilmington, MA). Libraries were gel purified, pooled, and sequenced to 100 nucleotides in the paired end mode on an Illumina HiSeq 2000 instrument. Read quality was assessed using FastQC (v 0.11.2; http://www.bioinformatics.babraham.ac.uk/projects/fastqc/) and adapter sequences were removed and the reads quality trimmed using Cutadapt software (v 1.4.1) (Martin 2011) using a quality cutoff 20 and a minimum length of 81. *De novo* genome assembly was performed using Velvet (v 1.2.10) (Zerbino and Birney 2008) using a k-mer length of 51. Contigs shorter than 1,000 bp were filtered out and the remaining contigs were used for downstream analyses.

The *G. sempervirens* assembled contigs were scaffolded using the synthetic long reads longer than 1499 bp and the SSPACE-LongRead tool (v1-1) (Boetzer and Pirovano 2014); the default alignment options and scaffolding options were used. Genome assembly quality was evaluated using CEGMA (Parra et al. 2007) and representation of RNA-sequencing reads (see below).

### *Gelsemium sempervirens* transcriptome analyses

For RNA-seq analysis, total RNA was extracted from five tissues (immature leaf, stem, stamens, pistils and petal) of *G. sempervirens* using the Qiagen RNeasy kit. RNA-seq libraries were constructed using the Kapa library preparation kit (Kapa Biosystems, Wilmington, MA) and sequenced on an Illumina HiSeq 2000 (100 bp, paired end). Read quality assessment, adapter removal, and quality trimming was performed as described for genomic DNA sequences. Cleaned RNA-seq reads were aligned to the *G. sempervirens* genome assemblies using TopHat (v 1.4.1) (Trapnell et al. 2009) in the single-end mode allowing a maximum of two mismatches; for reporting multiple mapping reads, up to 20 alignments were permitted. Alignment statistics were obtained using SAMtools (v 0.1.19) (Li et al. 2009).

### Human mRNA splicing analysis

The assembled long read sequences were processed to remove all poly-A reads, then aligned to hg19/GRCh37 with STAR version 2.3.1z (Dobin et al. 2013) and with GMAP version 2014-12-21(Wu and Watanabe 2005). BEDTools version 2.18.2 (Quinlan and Hall 2010) was used to count the number of reads mapped to each human gene, and Cufflinks version 2.1.1 (Trapnell et al. 2012) was used to quantify gene expression level. The gene annotations used in the analysis are from Ensemble (GRCh37.72). To identify splice junctions from the data, reads that uniquely aligned to the human genome were extracted, and RSEQC version 2.6.1 (Wang et al. 2012) was used to determine the locations where reads were split and to compare the resulting sets with known splice sites from GRCh37.72.

### Barcode fidelity determination in a plasmid mixture

Six plasmids were mixed, linearized by restriction enzyme digestion, and sequenced. Three of the six plasmids were BioBrick backbones differing only in their antibiotic resistance genes (pSB1C3, pSB1K3, and pSB1A3). The remaining three were pJexpress414 (DNA2.0, Menlo Park, CA) containing different variants of the *E. coli* outer membrane protein OmpA. Reads were sorted by barcode, and each bin of reads was searched for short sequences unique to each of the six plasmids. Unique sequence counts for the three OmpA plasmids were plotted for each barcode group in **Fig. S14**.

### Cleaning of spurious synthetic reads in *env* analysis

Because the *env* samples were sequenced to high coverage, reads containing sequencing errors in the barcode region were abundant enough that truncated synthetic reads were assembled from the reads associated with spurious barcodes. These synthetic reads were removed before further analysis by identifying barcodes with a Hamming distance from another barcode of one or two, and discarding the barcode with the shorter synthetic read. Surviving synthetic reads longer than 1.5 kb were aligned against the two known variant sequences using the EMBOSS (v 6.6.0) *water* local alignment software (Rice et al. 2000). Three *env*1/*env*2 chimeras were manually created by cutting and pasting together sequences from the two parents, and subjected to the same analysis.

### Code availability

Scripts used to assemble and analyze synthetic reads are available at https://github.com/jstapleton/synthetic_reads.

## Data Access

Raw sequencing data reported in this paper can be downloaded from the National Center for Biotechnology Information (NCBI) Sequence Read Archive, under the BioProject ID numbers PRJNA279503, PRJNA279434, PRJNA279490, PRJNA279203, PRJNA278980, PRJNA279403, PRJNA279401, and PRJNA279269. Assembled synthetic reads can be downloaded from figshare at http://dx.doi.org/10.6084/m9.figshare.1356287. Raw *G. sempervirens* whole-genome shotgun and RNA-seq reads have been deposited in the Sequence Read Archive under the BioProject number (###, to be made public upon publication). The scaffolded *G. sempervirens* assembly is available in the NCBI Whole Genome Shotgun Sequence database under accession number (###, to be made public upon publication). The Velvet and SSPACE assemblies of *G. sempervirens* are also available at the Data Dryad Digital Repository at the DOI (###, URL to be made public upon publication). Scripts used to assemble and analyze synthetic reads are available at https://github.com/jstapleton/synthetic_reads.

## Acknowledgments

We thank Richard Lenski for sharing the twenty-four experimentally evolved *E. coli* strains and we acknowledge the John Hannah chair endowment for funding their sequencing. We thank Jerry Dodgson and Hans Cheng for providing the *G. gallus* DNA sample, Irene Li and Aritro Nath for isolating mRNA samples, and the staff of the MSU RTSF Genomics Core for helpful suggestions. This research was supported by the National Institutes of Health under Ruth L. Kirschstein National Research Service Award F32GM099291 from the NIGMS to J.A.S, National Institutes of Health (5R21CA176854-02), MIIE Commercialization Grant (AGR 2014-0200), and MTRAC to T.A.W. Funds for the *G. sempervirens* genome and transcriptome analyses were provided by Michigan State University to C.R.B.

## Author Contributions

J.A.S. and T.A.W. conceived the technique. J.A.S. planned experiments, prepared samples for sequencing, wrote scripts to assemble synthetic reads, and analyzed data. T.A.W planned experiments and analyzed data. J.K. and C.R.B. isolated DNA from *Gelsemium* and *Solanum*. J.K. isolated RNA from *Gelsemium*. L.N. prepared *Gelsemium* Illumina libraries. J.K., J.H., and C.R.B. analyzed data. M.W. and C.C. provided mRNA and analyzed data. L.C.I. and C.T.B. analyzed *Gallus* data. R.M. provided DNA from evolved *E. coli*. B.B. and D.R.B. provided *env* samples and performed sequencing. J.A.S. and T.A.W. wrote the manuscript with input from the other authors.

## Disclosure Declaration

The authors declare competing financial interests in the form of a patent application covering aspects of the methods described in this manuscript.

## References

Acevedo A, Brodsky L, Andino R. 2014. Mutational and fitness landscapes of an RNA virus revealed through population sequencing. Nature 505: 686 – 690.

Bankevich A, Nurk S, Antipov D, Gurevich AA, Dvorkin M, Kulikov AS, Lesin VM, Nikolenko SI, Pham S, Prjibelski AD et al. 2012. SPAdes: a new genome assembly algorithm and its applications to single-cell sequencing. Journal of computational biology : a journal of computational molecular cell biology 19: 455 – 477.

Bentley DR Balasubramanian S Swerdlow HP Smith GP Milton J Brown CG Hall KP Evers DJ Barnes CL Bignell HR et al. 2008. Accurate whole human genome sequencing using reversible terminator chemistry. Nature 456: 53 – 59.

Boetzer M, Pirovano W. 2014. SSPACE-LongRead: scaffolding bacterial draft genomes using long read sequence information. BMC bioinformatics 15: 211.

Bolger AM, Lohse M, Usadel B. 2014. Trimmomatic: a flexible trimmer for Illumina sequence data. Bioinformatics 30: 2114 – 2120.

Branton D, Deamer DW, Marziali A, Bayley H, Benner SA, Butler T, Di Ventra M, Garaj S, Hibbs A, Huang XH et al. 2008. The potential and challenges of nanopore sequencing. Nature biotechnology 26: 1146 – 1153.

Burton DR, Desrosiers RC, Doms RW, Koff WC, Kwong PD, Moore JP, Nabel GJ, Sodroski J, Wilson IA, Wyatt RT. 2004. HIV vaccine design and the neutralizing antibody problem. Nature immunology 5: 233 – 236.

Consortium PGS, Xu X, Pan S, Cheng S, Zhang B, Mu D, Ni P, Zhang G, Yang S, Li R et al. 2011. Genome sequence and analysis of the tuber crop potato. Nature 475: 189 – 195.

Dobin A, Davis CA, Schlesinger F, Drenkow J, Zaleski C, Jha S, Batut P, Chaisson M, Gingeras TR. 2013. STAR: ultrafast universal RNA-seq aligner. Bioinformatics 29: 15 – 21.

Dunning AM, Talmud P, Humphries SE. 1988. Errors in the polymerase chain reaction. Nucleic Acids Res 16: 10393.

Georgiou G, Ippolito GC, Beausang J, Busse CE, Wardemann H, Quake SR. 2014. The promise and challenge of high-throughput sequencing of the antibody repertoire. Nature biotechnology 32: 158 – 168.

Hayashi K, Morooka N, Yamamoto Y, Fujita K, Isono K, Choi S, Ohtsubo E, Baba T, Wanner BL, Mori H et al. 2006. Highly accurate genome sequences of Escherichia coli K-12 strains MG1655 and W3110. Molecular systems biology 2: 2006.0007.

Hess M, Sczyrba A, Egan R, Kim TW, Chokhawala H, Schroth G, Luo SJ, Clark DS, Chen F, Zhang T et al. 2011. Metagenomic Discovery of Biomass-Degrading Genes and Genomes from Cow Rumen. Science 331: 463 – 467.

Hiatt JB, Patwardhan RP, Turner EH, Lee C, Shendure J. 2010. Parallel, tag-directed assembly of locally derived short sequence reads. Nature methods 7: 119 – 122.

Hong LZ, Hong S, Wong HT, Aw PP, Cheng Y, Wilm A, de Sessions PF, Lim SG, Nagarajan N, Hibberd ML et al. 2014. BAsE-Seq: a method for obtaining long viral haplotypes from short sequence reads. Genome biology 15: 517.

Jia JZ, Zhao SC, Kong XY, Li YR, Zhao GY, He WM, Appels R, Pfeifer M, Tao Y, Zhang XY et al. 2013. Aegilops tauschii draft genome sequence reveals a gene repertoire for wheat adaptation. Nature 496: 91 – 95.

Kuleshov V, Xie D, Chen R, Pushkarev D, Ma Z, Blauwkamp T, Kertesz M, Snyder M. 2014. Whole-genome haplotyping using long reads and statistical methods. Nature biotechnology 32: 261 – 266.

Li H. 2013. Aligning sequence reads, clone sequences and assembly contigs with BWA-MEM. arXiv:1303.3997v1302 [q-bio.GN]

Li H, Handsaker B, Wysoker A, Fennell T, Ruan J, Homer N, Marth G, Abecasis G, Durbin R. 2009. The Sequence Alignment/Map format and SAMtools. Bioinformatics 25: 2078 – 2079.

Lundin S, Gruselius J, Nystedt B, Lexow P, Kaller M, Lundeberg J. 2013. Hierarchical molecular tagging to resolve long continuous sequences by massively parallel sequencing. Scientific reports 3: 1186.

Magoc T, Salzberg SL. 2011. FLASH: fast length adjustment of short reads to improve genome assemblies. Bioinformatics 27: 2957 – 2963.

Margulies M, Egholm M, Altman WE, Attiya S, Bader JS, Bemben LA, Berka J, Braverman MS, Chen YJ, Chen ZT et al. 2005. Genome sequencing in microfabricated high-density picolitre reactors. Nature 437: 376 – 380.

Martin M. 2011. Cutadapt removes adapter sequences from high-throughput sequencing reads. 2011 17: 10 – 12.

McCoy RC, Taylor RW, Blauwkamp TA, Kelley JL, Kertesz M, Pushkarev D, Petrov DA, Fiston-Lavier AS. 2014. Illumina TruSeq Synthetic Long-Reads Empower De Novo Assembly and Resolve Complex, Highly-Repetitive Transposable Elements. PloS one 9: e106689.

Menon R, Im H, Zhang EY, Wu SL, Chen R, Snyder M, Hancock WS, Omenn GS. 2014. Distinct splice variants and pathway enrichment in the cell-line models of aggressive human breast cancer subtypes. Journal of proteome research 13: 212 – 227.

Metzker ML. 2010. Applications of Next-Generation Sequencing Sequencing Technologies - the Next Generation. Nat Rev Genet 11: 31 – 46.

Miller MR, Dunham JP, Amores A, Cresko WA, Johnson EA. 2007. Rapid and cost-effective polymorphism identification and genotyping using restriction site associated DNA (RAD) markers. Genome research 17: 240 – 248.

Parra G, Bradnam K, Korf I. 2007. CEGMA: a pipeline to accurately annotate core genes in eukaryotic genomes. Bioinformatics 23: 1061 – 1067.

Picelli S, Bjorklund AK, Faridani OR, Sagasser S, Winberg G, Sandberg R. 2013. Smart-seq2 for sensitive full-length transcriptome profiling in single cells. Nature methods 10: 1096 – 1098.

Picelli S, Faridani OR, Bjorklund AK, Winberg G, Sagasser S, Sandberg R. 2014. Full-length RNA-seq from single cells using Smart-seq2. Nature protocols 9: 171 – 181.

Quinlan AR, Hall IM. 2010. BEDTools: a flexible suite of utilities for comparing genomic features. Bioinformatics 26: 841 – 842.

Rice P, Longden I, Bleasby A. 2000. EMBOSS: the European Molecular Biology Open Software Suite. Trends in genetics : TIG 16: 276 – 277.

Rubin CJ, Zody MC, Eriksson J, Meadows JR, Sherwood E, Webster MT, Jiang L, Ingman M, Sharpe T, Ka S et al. 2010. Whole-genome resequencing reveals loci under selection during chicken domestication. Nature 464: 587 – 591.

Rungpragayphan S, Kawarasaki Y, Imaeda T, Kohda K, Nakano H, Yamane T. 2002. High-throughput, Cloning-independent Protein Library Construction by Combining Single-molecule DNA Amplification with in Vitro Expression. Journal of molecular biology 318: 395 – 405.

Saghai-Maroof MA, Soliman KM, Jorgensen RA, Allard RW. 1984. Ribosomal DNA spacer-length polymorphisms in barley: mendelian inheritance, chromosomal location, and population dynamics. Proceedings of the National Academy of Sciences of the United States of America 81: 8014 – 8018.

Sharma SK, Bolser D, de Boer J, Sonderkaer M, Amoros W, Carboni MF, d’Ambrosio JM, de la Cruz G, Di Genova A, Douches DS et al. 2013. Construction of reference chromosome-scale pseudomolecules for potato: integrating the potato genome with genetic and physical maps. G3 (Bethesda, Md) 3: 2031 – 2047.

Sharon I, Kertesz M, Hug LA, Pushkarev D, Blauwkamp TA, Castelle CJ, Amirebrahimi M, Thomas BC, Burstein D, Tringe SG et al. 2015. Accurate, multi-kb reads resolve complex populations and detect rare microorganisms. Genome research doi:10.1101/gr.183012.114.

Souza V, Turner PE, Re L. 1997. Long-term experimental evolution in Escherichia coli. V. Effects of recombination with immigrant genotypes on the rate of bacterial evolution. Journal of Evolutionary Biology 10: 743 – 769.

Stapleton JA, Swartz JR. 2010. A cell-free microtiter plate screen for improved [FeFe] hydrogenases. PloS one 5: e10554.

Trapnell C, Pachter L, Salzberg SL. 2009. TopHat: discovering splice junctions with RNA-Seq. Bioinformatics 25: 1105 – 1111.

Trapnell C, Roberts A, Goff L, Pertea G, Kim D, Kelley DR, Pimentel H, Salzberg SL, Rinn JL, Pachter L. 2012. Differential gene and transcript expression analysis of RNA-seq experiments with TopHat and Cufflinks. Nature protocols 7: 562 – 578.

Voskoboynik A, Neff NF, Sahoo D, Newman AM, Pushkarev D, Koh W, Passarelli B, Fan HC, Mantalas GL, Palmeri KJ et al. 2013. The genome sequence of the colonial chordate, Botryllus schlosseri. eLife 2: e00569.

Wang L, Wang S, Li W. 2012. RSeQC: quality control of RNA-seq experiments. Bioinformatics 28: 2184 – 2185.

Wu NC, De La Cruz J, Al-Mawsawi LQ, Olson CA, Qi H, Luan HH, Nguyen N, Du Y, Le S, Wu TT et al. 2014. HIV-1 quasispecies delineation by tag linkage deep sequencing. PloS one 9: e97505.

Wu TD, Watanabe CK. 2005. GMAP: a genomic mapping and alignment program for mRNA and EST sequences. Bioinformatics 21: 1859 – 1875.

Zerbino DR, Birney E. 2008. Velvet: algorithms for de novo short read assembly using de Bruijn graphs. Genome research 18: 821 – 829.

